# Functional Characterization of a Lassa Virus Fusion Inhibitors Adaptive Mutant

**DOI:** 10.1101/2020.12.23.424274

**Authors:** Jiao Guo, Guangshun Zhang, Yang Liu, Junyuan Cao, Mengmeng Zhang, Xiaohao Lan, Yueli Zhang, Chenchen Liu, Gengfu Xiao, Wei Wang

## Abstract

Lassa virus (LASV) glycoprotein complex (GPC) contains retained stable-signal peptide (SSP), GP1, and GP2. SSP interacts with GP2 and provides an interface targeted by numerous fusion inhibitors. Serially passaging of LASV with inhibitors allowed some adaptive mutants to be obtained of which most had mutations located in the transmembrane (TM) domain of GP2. In the current study, we focused on the F446L mutant, which is reported to confer resistance to ST-series inhibitors. We found that F446L conferred cross-resistance to structurally distinct inhibitors. Furthermore, F446L increased the fusion activities of LASV and Mopeia virus GPC, elevating the pH threshold for fusion of LASV and promoting fusion of MOPV at neutral pH. F446L exerted little effect on the pseudotype viral growth profile or thermostability. By introducing other residues to the conserved F446 locus, it was found that this site was less compatible with a similar tyrosine residue and was intolerable to charged residues. These results help characterize the fusion inhibitor target located in the TM domain of GP2, which should be useful for drug and vaccine design.

**IMPORTANCE:** The LASV SSP-GP2 interface provides an Achilles heel that is targeted by numerous inhibitors. However, the emergence of resistant viruses is a major concern for direct antiviral drugs. In this study, we investigated the F446L mutant located in the GPC TM domain to determine the relationship between drug resistance, membrane fusion activity, viral growth kinetics, and thermostability. These results will be helpful in monitoring drug-resistant variants, as well as the advancement of drug and vaccine design.

## INTRODUCTION

Lassa virus (LASV) is an enveloped virus with a bi-segmented, negative-sense RNA genome that belongs to the genus *Mammarenavirus* (family *Arenaviridae*). Mammarenaviruses include 39 species, which are divided into New World (NW) and Old World (OW) viruses based on virus genetics, serology, antigenic properties, and geographical relationships (1, 2). The OW viruses LASV and Lujo virus (LUJV), and NW viruses Junín virus (JUNV), Machupo virus (MACV), Guanarito virus (GTOV), Chapare virus (CHAPV), and Sabiá virus (SBAV) can cause severe hemorrhagic fever and are categorized as biosafety level (BSL) 4 pathogens (3, 4).

The bi-segmented RNA genomes contain the small (S) and large (L) segments. The S segment encodes the glycoprotein complex (GPC) and nucleoprotein, while the L segment encodes the matrix protein (Z) and viral polymerase. The mature GPC is present on the surface of the virion and consists of three subunits, the stable-signal peptide (SSP), receptor-binding subunit GP1, and membrane fusion subunit GP2 (5–7). The three subunits are non-covalently bound and form a (SSP/GP1/GP2)_3_ trimeric complex (Fig. 1A and 1B). Notably, the unusual retained SSP plays an essential role in GPC-mediated membrane fusion and cell entry. SSP anchors to the viral membrane by interacting with the membrane proximal external region, the transmembrane domain (TM), and the cytoplasmic tail of GP2, stabilizing the pre-fusion conformation of GPC and providing an interface that is targeted by fusion inhibitors (5–7). It has been widely reported that mammarenaviruses generate adaptive mutants against entry inhibitors mostly through mutations located at the SSP-GP2 interface.

**Fig. 1.**
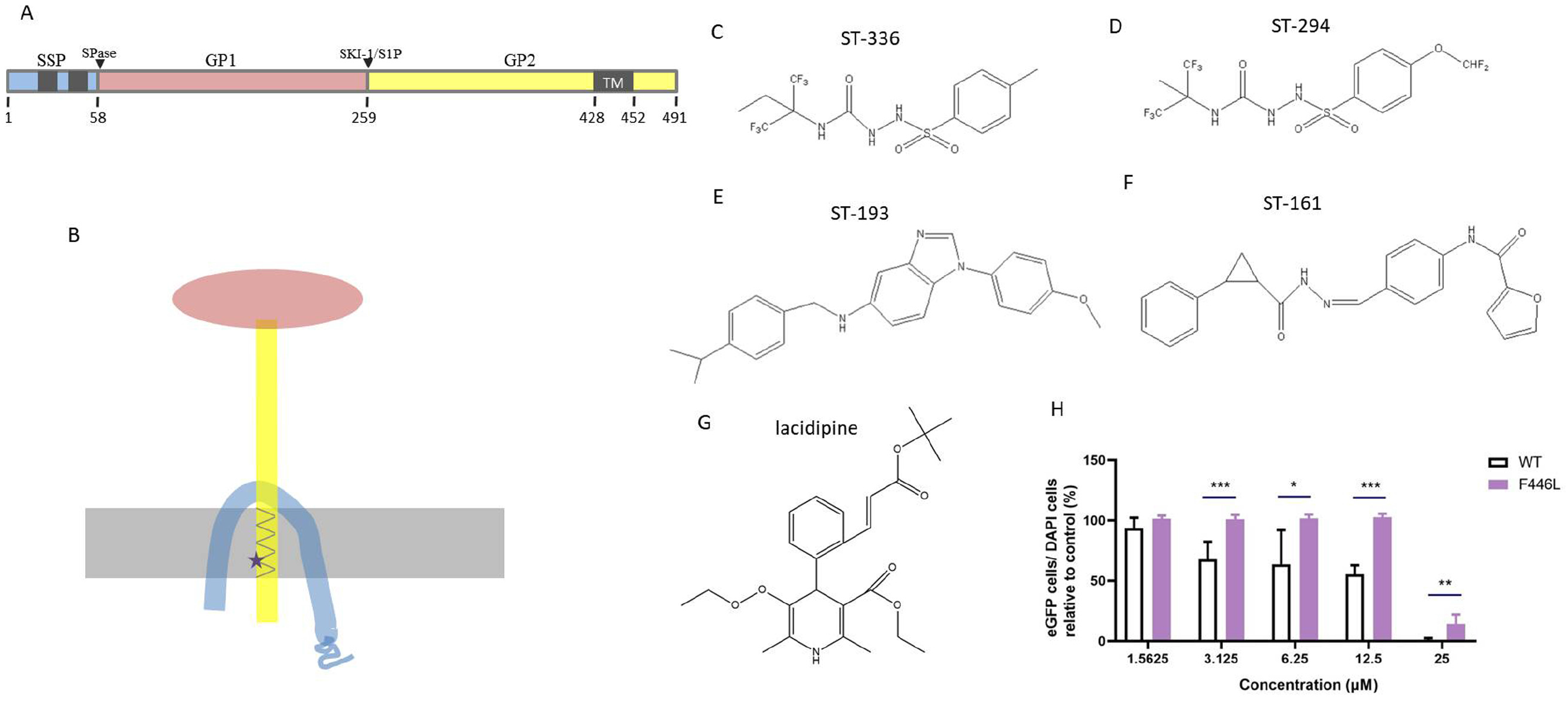
LASV F446L conferred resistance to distinct inhibitors. (**A**) Schematic diagram of the LASV GPC open reading frame. TM domains in SSP and GP2 are shown in light gray. F446 is labeled with an asterisk. The drawing is not to scale. (**B**) Schematic diagram of the SSP/GP1/GP2 heterotrimer. Structure of ST-336 (**C**), ST-294 (**D**), ST-193 (**E**), ST-161 (**F**), and lacidipine (**G**). (**H**) The F446L mutation conferred resistance to lacidipine. Vero cells were incubated with lacidipine at the indicated concentration; 1 h later, LASV_WTrv_ and LASV_F446Lrv_ were added to the cells (MOI: 0.1) and adsorbed for 1 h. The eGFP signal was detected 23 h later using an Operetta High-Content Imaging System. Data are presented as means ± SDs from 4–6 independent experiments. *P < 0.05, ** P < 0.01, *** P < 0.001.

Intriguingly, some adaptive mutants show cross-resistance to other entry inhibitors (8). The GPC F446L mutant of LASV and corresponding mutants in other mammarenaviruses, such as JUNV F438L, have been identified among these adaptive mutants to be resistant to multiple small-molecule inhibitors (8–11). To date, insight into the mechanisms underlying resistance is limited. Whether the adaptive mutant de-stabilizes the prefusion conformation of GPC or if the native residue of the adaptive mutant serves as a viral target that interacts directly with the inhibitors is unclear.

In the current work, we aimed to characterize the effects of the F446L mutant on GPC-mediated membrane fusion, receptor binding, virus thermostability, growth kinetics, and fitness. The results should provide important context for investigating the relationship between viral variants and their virulence, which will contribute to the advancement of drug and vaccine design.

## RESULTS

### LASV GPC F446L exhibited resistance to structurally distinct inhibitors

As noted, the F446L mutation is reported to confer resistance to numerous inhibitors. Passaging LASV with ST-161 derivative led to the isolation of the F446L mutant, while passaging Tacaribe virus (TCRV) in the presence of ST-336 and ST-193 led to the F436I mutant, corresponding to LASV GPC F446 mutants (9–11). The F446L mutation has been reported to confer resistance to these ST-series inhibitors and their structurally similar inhibitors (Fig. 1C–1F) (8, 9). This raises the question whether F446L confers resistance to other mammarenavirus entry inhibitors that caused adaptive mutants with changes at loci other than the F446L change itself. To this end, we investigated the sensitivity of LASV_F446Lrv_ to lacidipine, in accordance with our previous reported that lacidipine caused the adaptive T40K mutant in LASV GPC SSP (6). As shown in Fig. 1H, lacidipine showed a dose-dependent inhibition against LASV_WTrv_ infection. Lacidipine (25 μM) almost abolished the LASV_WTrv_ infection, which was consistent with our previous report (6). Meanwhile, LASV_F446Lrv_ exhibited robust resistance to lacidipine. LASV_F446Lrv_ was able to establish approximately 100% infection, even after treatment with 12.5 μM lacidipine. When treated with 25 μM lacidipine, LASV_F446Lrv_ was capable of establishing approximately 15% infection, which was significantly higher than that of wild-type (WT) LASV (P = 0.00359).

### LASV GPC F446L promoted efficient fusion

F446 is located in the transmembrane domain of GP2 (Fig 1A and 1B). As GP2 is the fusion subunit of the glycoprotein and TM plays an essential role in the GPC-mediated fusion activity (9), we further evaluated the effect of F446L on GPC-mediated fusion. We transfected 293t cells with WT or F446L GPC and 24 h later treated the cells with buffers of various pH for 15 min. Syncytium formation was then observed 4 h later. As shown in Fig. 2A, both WT and F446L GP2 could lead to a complete fusion when treated with buffer at either pH 4.5 or 5.0. In comparison, WT GP2 was able to induce only partial syncytium formation after exposure to pH 5.5, while F446L still demonstrated complete fusion. To confirm this finding, a quantitative evaluation was performed using a dual-luciferase assay. Consistent with the results from the qualitative assay, the quantitative assay indicated that F446L was able to reach a much greater degree of fusion activity compared to that of WT when treated with either pH 5.5 or pH 6.0 buffer (Fig. 2B). As pH 6.0 and pH 7.0 were barely able to trigger GPC-mediated fusion for either WT or F446L, the qualitative assay was not conducted at these two pH values. Based on our findings that F446L raised the pH threshold for GPC-mediated fusion, we investigated whether F446L entered the cells through the pH-dependent endocytic pathway. It was found that LASV_F446Lrv_ remained sensitive to NH_4_Cl treatment, similar to that of LASV_WTrv_ and VSV_rv_, which indicated the entry of LASV_F446Lrv_ remained pH-dependent (Fig. 2C).

**Fig. 2.**
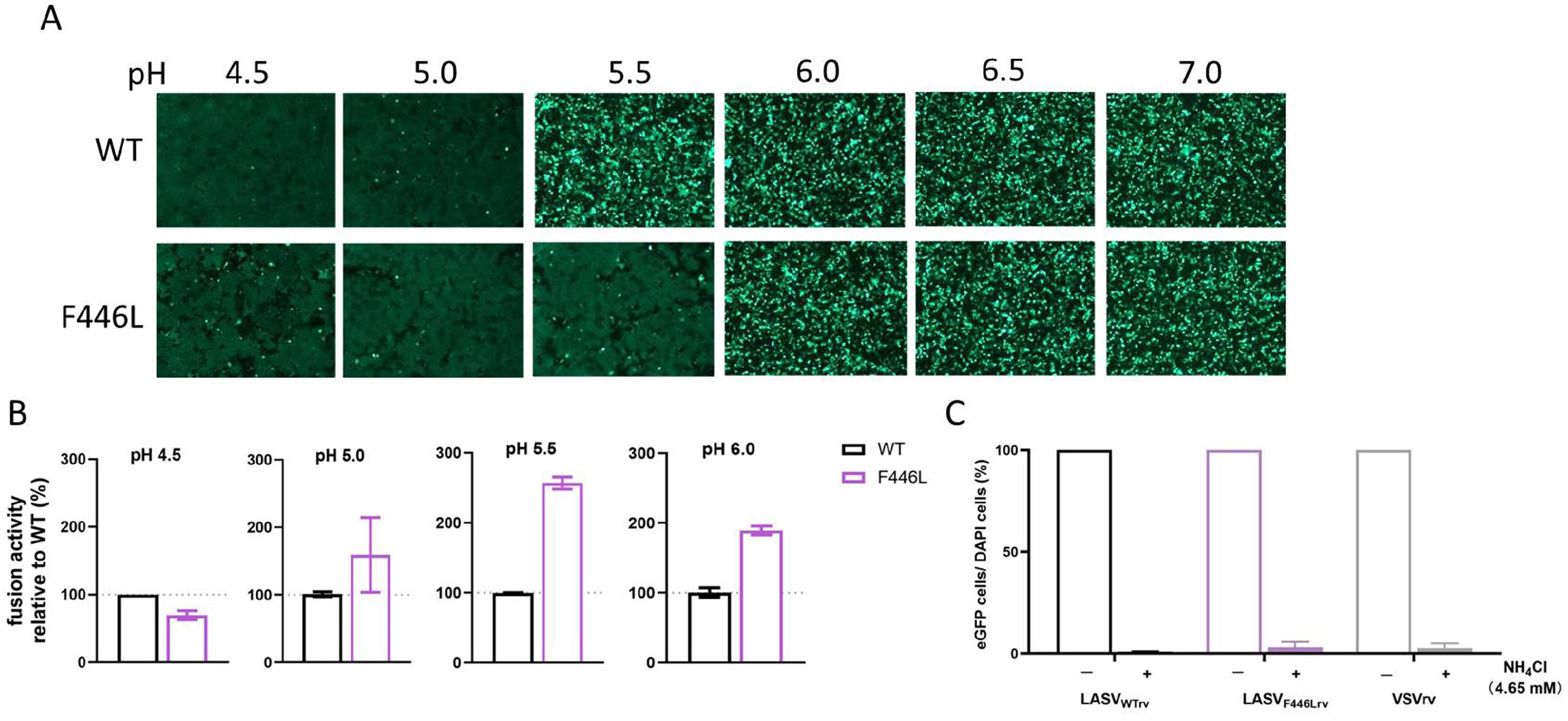
Membrane fusion activity of the F446L mutant. (**A**) Qualitative evaluation of WT and F446L fusion activities. The 293T cells were transfected with WT or F446L GPC; 24 h later, the cells were treated with acidified buffer (pH 4.5 to 7) for 15 min. The cells were then placed in DMEM at a neutral pH. Syncytium formation was visualized 1 h later using fluorescent microcopy. Images are representative fields from three independent experiments. (**B**) Quantitative evaluation of LASV WT and F446L fusion activities. The 293T cells transfected with both GPC and pCAG-T7 then co-cultured at a ratio of 3:1 with cells transfected with pT7EMCVLuc and the pRL-CMV control vector. Fusion activity was evaluated by measuring Fluc activity, which was standardized to Rluc activity. Data are presented as means ± SDs from three independent experiments. *** P < 0.001.

### F446L retained Alfa-dystroglycan (α-DG) binding ability

Alfa-DG is the primary receptor of LASV (12, 13). It was expected that a TM mutant would have little effect on receptor binding of the glycoprotein. To investigate the binding ability of F446L, A549 cells were inoculated with LASV_WTrv_ and LASV_F446Lrv_ at a multiplicity of infection (MOI) of 0.1, adsorbed for 1 h, and the eGFP-positive cells counted 23 h later. As a control, we used the monoclonal antibody (MAb) IIH6, which recognizes a functional glycan epitope on α-DG and thus competes with LASV binding (14, 15). As shown in Fig. 3, there were 1420 ± 359 eGFP-positive A549 cells/96-well plate in the LASV_WTrv_ infected group, while the number of dropped to approximately 388 when treated with IIH6. Similarly, there were 1169 ± 322 positive cells/96-well plate in the LASV_F446Lrv_ group without treatment with IIH6 and the number dropped to approximately 301 after IIH6 treatment. There was no significant difference in the number of eGFP-positive cells between LASV_WTrv_ and LASV_F446Lrv_, either in the absence (P = 0.0729) or presence (P = 0.151) of IIH6, indicating LASV_F446Lrv_ retained its ability to bind the primary receptor on the cell surface.

**Fig. 3.**
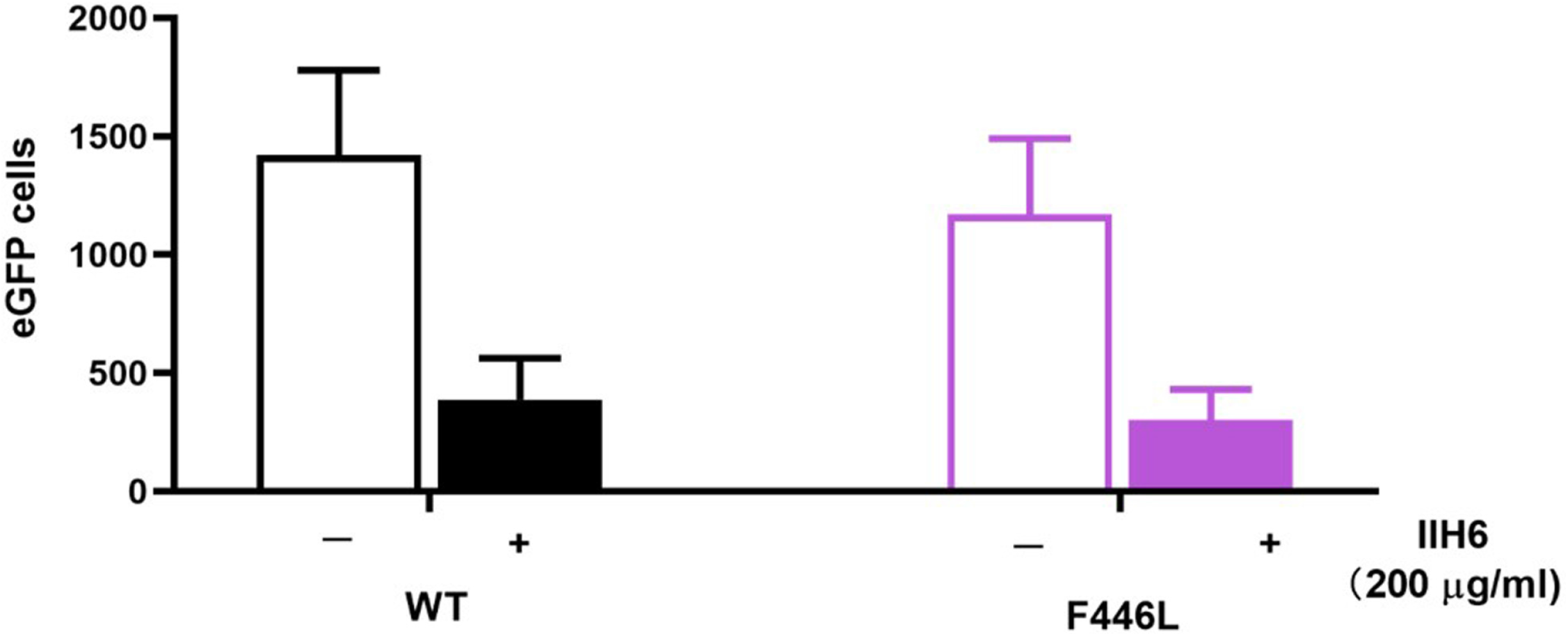
LASV_F446Lrv_ continued to utilize α-DG as the primary receptor. A549 cells were incubated with monoclonal antibody IIH6 (200 μg/ml) at 37°C for 1 h. LASV_WTrv_ and LASV_F446Lrv_ were added (MOI, 0.1) and incubated at 4℃ for an additional 1 h. The eGFP-positive cells were counted using an Operetta High-Content Imaging System 24 h later. Data are presented as means ± SDs from 2–3 independent experiments.

### LASV_F446Lrv_ possessed thermostability similar to LASV_WTrv_

As F446L may alter the fusion activity of GPC, we investigated whether F446L impaired the stability of the LASV GPC. LASV_F446Lrv_, in parallel with LASV_WTrv_, was incubated at various temperatures between 25°C and 55°C and then inoculated onto Vero cells. The eGFP-positive cells were analyzed using a high-content imaging system. The results showed the transduction activities of both LASV_F446Lrv_ and LASV_WTrv_ decreased as the temperature increased. A dramatic decrease occurred at 47°C and both viruses lost transduction activities at 55°C. Notably, LASV_F446Lrv_ exhibited a similar thermal inactivation profile as that of LASV_WTrv_ (Fig. 4A). Likewise, when both viruses were incubated at 37°C and 25°C, the transduction activities decreased over time, but with no differences between LASV_F446Lrv_ and LASV_WTrv_ (Fig. 4B and 4C). These results suggested the contribution of F446L to the increase in fusogenic activity was not related to the thermostability of the recombinant virus.

**Fig. 4.**
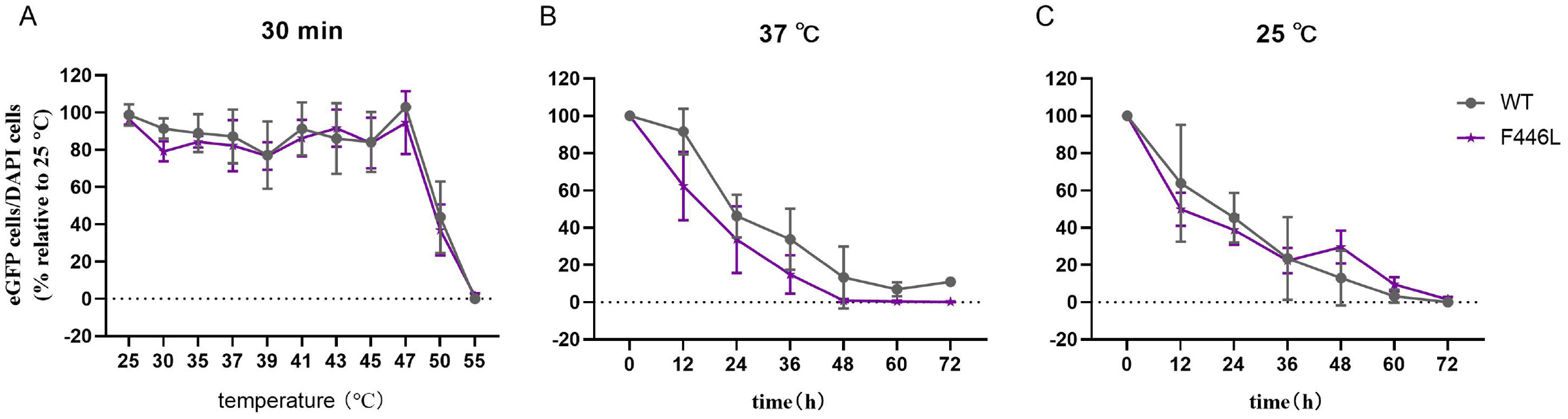
Thermostability of LASV_WTrv_ and LASV_F446Lrv_. LASV_WTrv_ and LASV_F446Lrv_ were incubated at the indicated temperature (25°C to 55°C) for 30 min (**A**), or for the indicated time (0 to 72 h) at 37°C (**B**) or 25°C (**C**) and then plaque assayed using Vero cells. Data are presented as means ± SDs for four independent experiments.

### Residue 446 in TM domain regulated virus growth kinetics

As F446L did not alter the thermostability of the virus, we compared the growth kinetics of LASV_F446Lrv_ and LASV_WTrv_. LASV_F446Lrv_ exhibited a similar multiple steps growth curve to that of LASV_WTrv_ during the first 72 h when Vero cells were inoculated at an MOI of 0.01. Both viruses reached maximum titers at 24 h post-infection (p.i.) with similar titers of approximately 10^8^ plaque forming units (PFU)/mL. However, the viral titer of LASV_F446Lrv_ decreased dramatically after 72 h, while the titer of LASV_WTrv_ began to decrease starting at 120 h (Fig. 5).

**Fig. 5.**
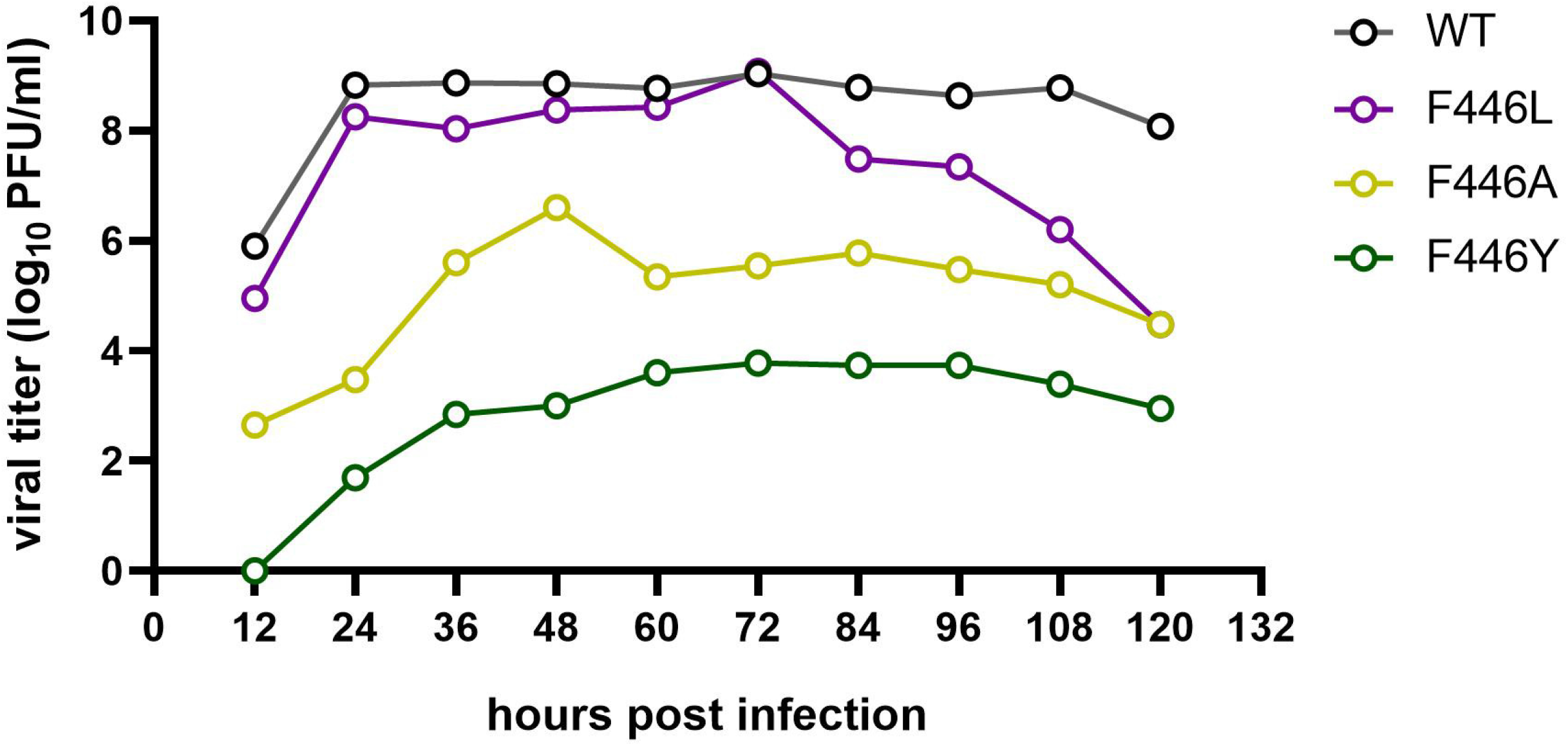
Recombinant viruses multi-step growth curves. Vero cells were infected with WT or mutant recombinant viruses at an MOI of 0.01 and allowed to adsorb for 1 h. Supernatant samples were collected at indicated time points post infection and plaque assayed for virus titer.

To investigate whether residue 446 in TM domain was tolerable for other residue changes, we introduced alanine (A), a positively charged arginine (R), a negatively charged aspartic acid (D), and tyrosine (Y) with its benzene ring at this site and evaluated the fitness of the mutant recombinant viruses. Intriguingly, stable infectious viral particles could not be isolated for either the F446R or F446D mutants, suggesting charged residues were incompatible with this site. F446A and F446Y mutants could generate infectious viral particles, but the viral titers were much lower for both recombinant viruses compared to those of WT. Notably, the F446Y mutant, with a structure similar to that of WT at position 446 in TM domain, generated a viral titer of 10^3^–10^4^ PFU/mL.

### Fusion occurred at neutral pH for MOPV with the corresponding F444L mutation

As TM domain is highly conserved in mammarenavirus (Fig. 6A), we introduced corresponding F446 mutations in other mammarenavirus GPCs and investigated the effects on GPC-mediated fusion. Unexpectedly, we found that the corresponding F444L mutation in MOPV could lead to membrane fusion, even in the absence of acidic conditions. At 16 h p.i. infection with MOPV GPC F446L, definite syncytium formation was observed in 293 cells with the results being similar to that of cells infected with the MOPV GPC WT virus treated with pH 5.5 (Fig. 6B). We also introduced the corresponding mutation into other mammarenaviruses and evaluated whether the mutation would increase the pH threshold for GPC-mediated fusion. As shown in Fig. 6C, the corresponding mutants of LCMV, JUNV, and GTOV exhibited decreased GPC-mediated fusion activities, while the substitutions had little effect on the MACV, SBAV, and CHAPV mutants. These results indicated the effects of this mutation on increasing fusogenicity were largely limited to LASV and was most closely related to MOPV.

**Fig. 6.**
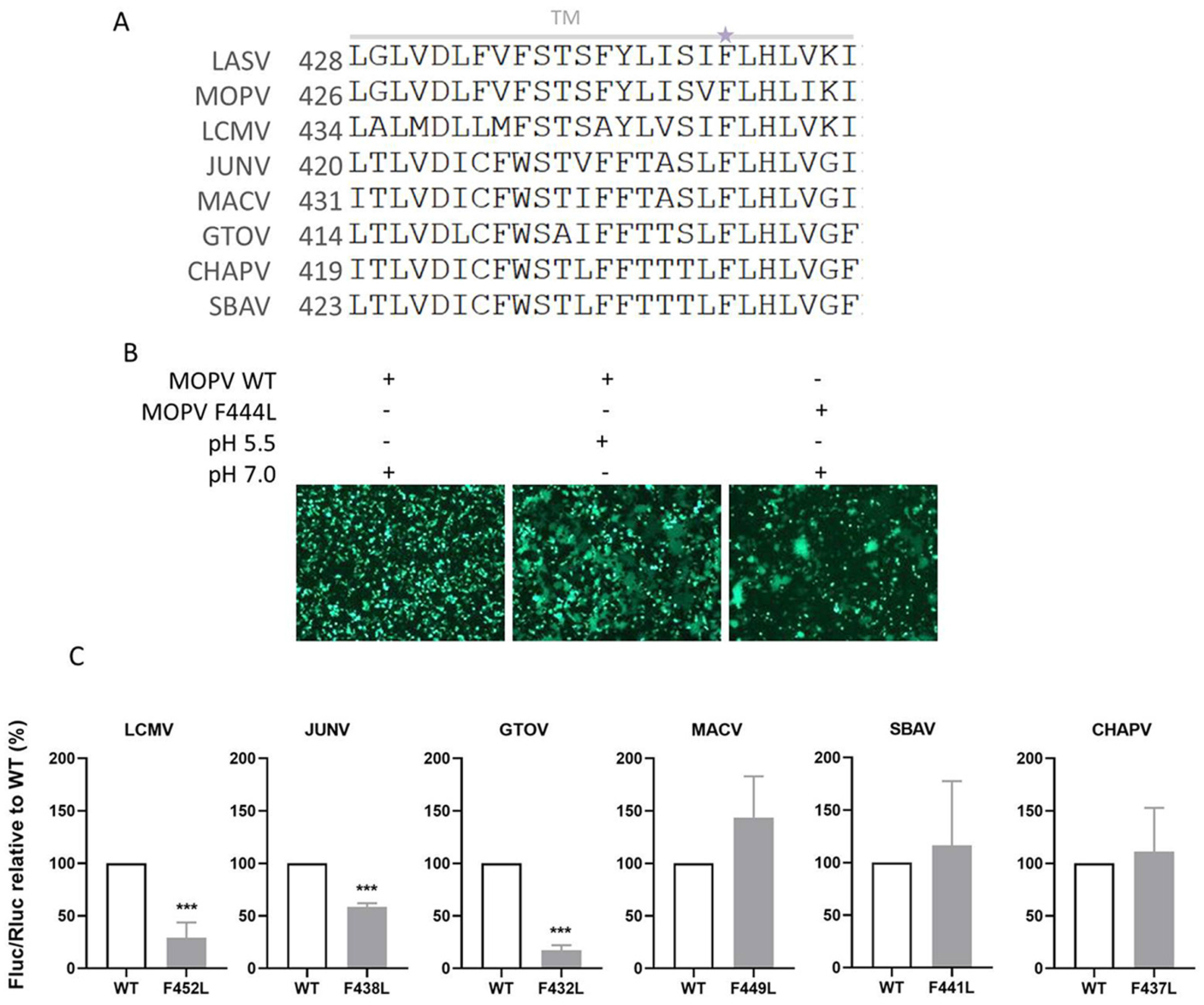
Effects of corresponding mutations in other mammarenavirus on GPC-mediated fusion activity. (**A**) Sequence alignment of the TM domain of mammarenavirus GPCs. LASS F446 is highlighted with a purple asterisk. (**B**) MOPV corresponding F444L mutation increased virus fusion activity. The 293T cells were transfected with MOPV WT or F444L GPC; 24 h later, the cells were treated with acidic or neutral buffer for 15 min. Syncytium formation was visualized 1 h later using fluorescent microcopy. Images are representative fields from five independent experiments. (**C**) Quantitative evaluation of the effects of corresponding mutations on other mammarenavirus GPC-mediated fusion activities. The 293T cells were transfected with both GPC (WT or mutant) and pCAG-T7 and then co-cultured at a ratio of 3:1 with cells transfected with pT7EMCVLuc and the pRL-CMV control vector. Fusion activity was evaluated by measuring Fluc activity, which was standardized to Rluc activity. Data are presented as means ± SDs for 3–6 independent experiments. *** P < 0.001

## DISCUSSION

The unique retention of SSP and its interaction with GP2 provides an interface target for numerous structurally distinct inhibitors (5–7). However, considering the high mutation rate of arenaviruses, drug resistance is a constant major concern in the context of direct antiviral agents. Among the reported drug-resistant variants, LASV F446L and its corresponding mutants in other arenaviruses have been extensively recorded. In the current study, we found that F446L conferred cross-resistance, not only to the ST series of inhibitors, but also to lacidipine, which we previously reported resulted in the emergence of an adaptive mutant with a T40K change in SSP (6). The detailed molecular mechanism underlying drug resistance is unknown. Being embedded in the TM domain of GP2, the natural phenylalanine residue at this site may be accessible to highly lipophilic fusion inhibitors. On the other hand, the change from phenylalanine to leucine may directly or allosterically prevent the inhibitor-binding or to modify the effect of binding, or contribute to an increase in virulence and thereby make the inhibitor ineffective.

Mutations in glycoproteins identified during pandemic outbreaks are usually accompanied by enhanced pathogenesis and transmission. The spike protein mutant D614G that has emerged during the expanding Coronavirus Disease 2019 (COVID-19) pandemic has been reported to increase the infectivity and transmission of SARS-CoV-2, the etiological virus (16–18). Similarly, the Ebola glycoprotein A82V mutant in the 2013–2016 epidemics increased infectivity in human cells and the increase in infectivity was accompanied by increased fusion activity (19, 20). Accordingly, we evaluated the sequences of clinical isolates reported for the 2018 LASV outbreak in Nigeria (21) and found that the F446L mutant was absent. On the other hand, glycoprotein mutants that emerge during serial passaging for vaccine development are always accompanied by attenuated virulence. The only currently approved arenavirus vaccine strain for JUNV, Candid #1, possesses six amino acid substitutions with the major determinant of attenuation being GPC F427I, which is located in the TM domain (22–24). Notably, corresponding to the JUNV F427I substitution, MACV F438I and TCRV F425L resulted in significant reduction of virulence in mouse models compared to that of the WT parent viruses. The corresponding LASV V434I mutant promotes fusion activity at neutral pH and also confers resistance to a fusion inhibitor (23–25). In the current study, we found that LASV F446L also increased fusion activity and the corresponding MOPV F444L mutation led to membrane fusion at neutral pH. Meanwhile, F446L had little effect on the growth kinetics of LASV, which was consistent with our previous report that adaptive mutants generally do not impair virus fitness (6, 26). However, this position in the TM domain of GPC only tolerated phenylalanine and leucine, not charged residues or tyrosine with the most structural similarity, suggesting the highly conserved phenylalanine is nearly indispensable for virion assembly and GPC function. Typically, glycoprotein mutations increase fusion activity at the cost of virus stability (20, 27–30). In the current study, the F446L mutant showed little effect on the thermostability of LASVrv, suggesting there may be a delicate gap between energy thresholds of fusion and thermostability; GPC_F446L_ might maintain a metastable state. We sequenced the LASV_F446Lrv_ genome after more than eight serial passages and observed no additional mutations or any reversion mutations, indicating that L446 could generate stable progeny virions, even in the absence of inhibitor selection pressure. Importantly, whether F446L in native LASV exhibits similar thermostability and whether LASV_F446Lrv_ is less virulent during infection *in vivo* need to be further investigated.

## MATERIALS AND METHODS

### Cells, plasmids, and viruses

BHK-21, HEK 293T, Vero, and A549 cells were cultured in Dulbecco’s modified Eagle’s medium (DMEM; HyClone, Logan, UT, USA) supplemented with 10% fetal bovine serum (FBS; Gibco, Grand Island, NY, USA). The GPC genes of LASV (Josiah strain, GenBank HQ688673.1), MOPV (GenBank AY772170.1), LCMV (Armstrong strain, GenBank AY847350.1), GTOV (GenBank NC_005077.1), JUNV (XJ13 strain, GenBank NC_005081.1), MACV (Carvallo strain, GenBank NC_005078.1), SBAV (GenBank U41071.1), and CHAPV (GenBank NC_010562.1) were synthesized (Sangon Biotech, Shanghai, China) and subcloned into pCAGGS vector. The primers used for generating the point mutation are listed in Table 1.

**Table 1.**
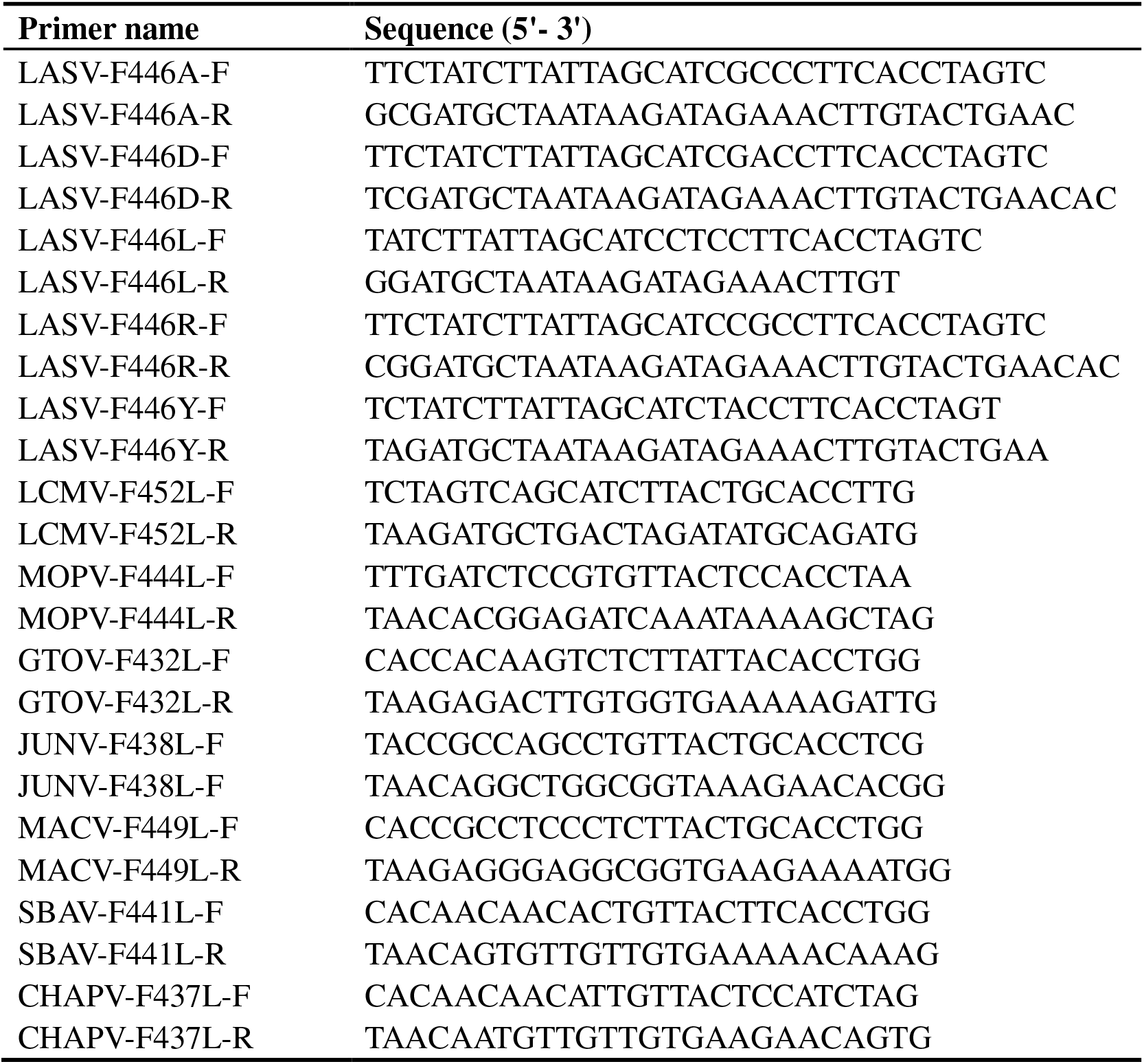
Primers for point mutations.

Plasmids encoding each protein component of the vesicular stomatitis virus (VSV), pBS-N, pBS-P, pBS-L, and pBS-G, were purchased from Kerafast. Plasmid pVSV∆G-eGFP (Plasmid #31842, Addgene) was modified to construct pVSV∆G-eGFP-GPC, which was used to generate LASVrv. Briefly, LASVrv virus was produced by transfecting BHK-21 cells in 6-wells plates that had been previously infected at an MOI of 5 with a recombinant vaccinia virus (vTF7-3) encoding T7 RNA polymerase. After a 45-min adsorption period, the infected cells were transfected with 11 μg of a mixture of plasmids including pVSV∆G-eGFP-GPC, pBS-N, pBS-P, pBS-G, and pBS-L at a 5:3:5:8:1 ratio (31, 32). At 48 h post transfection, the supernatants were collected and filtered to remove the vaccinia virus and then used to inoculate the BHK-21 cells that had been transfected 24 h previously with pCAGGS-VSV-G. The supernatant was collected 36 h later, filtered, aliquoted, and frozen at −80°C until further use.

### Membrane fusion assay

For qualitative evaluation of membrane fusion activity, 293T cells co-transfected with WT or mutant pCAGGS-GPC and pEGFP-N1 were incubated for 15 min with citric acid and sodium citrate buffers at the indicated pH. Syncytium formation was visualized 1 h later. For quantitative evaluation of membrane fusion activity, 293T cells in 24-well plates were transfected with pCAGGS-GPC (0.25 μg/well) and pCAGT7 (0.25 μg/well). Additional 293T cells in 6-well plates were transfected with pT7EMCVLuc (2.5 μg/well) and pRL-CMV (0.1 μg/well). The transfected cells were then co-cultured at a ratio of 3:1. The plasmids used in the assay were kindly provided by Yoshiharu Matsuura (Osaka University, Osaka, Japan). Twelve hours after co-culturing, the cells were treated with acidic buffer for 15 min and then continued to be incubated. Luciferase activities were measured 24 h later using a Dual-Glo Luciferase Assay (Promega, Madison, WI, USA) (33–35). Cell fusion activity was determined according to firefly luciferase (Fluc) activity standardized to renilla luciferase (Rluc) activity.

### IIH6 inhibition assay

A549 cells were pretreated with IIH6 (sc-53987; Santa Cruz Biotechnology, Santa Cruz, CA, USA) at 37°C for 1 h (14, 15). The cells were then incubated with LASV_WTrv_ or LASV_F446Lrv_ (MOI: 0.1) at 4°C for 1 h. After being extensively washed with phosphate-buffered saline (PBS), the cells were incubated at 37°C for an additional 24 h. The eGFP-positive cells were counted using an Operetta High-Content Imaging System (Perkin Elmer).

### Thermostability assay

LASV_WTrv_ or LASV_F446Lrv_ were resuspended in DMEM in the absence of serum and incubated for the indicated time at 37°C or 25°C, or incubated for 30 min at the indicated temperature (20, 27). The viruses were then used to infect Vero cells, which were evaluated for infection 23 h later using the Operetta High-Content Imaging System.

### Virus growth kinetics

Vero cells in 6-well plates were infected with WT or mutant recombinant LASV at an MOI of 0.01 and allowed to adsorb for 1 h at 37°C. The infected cells were then incubated and at each indicated time point, 200 μL of tissue culture supernatant was removed for analysis by the plaque assay. After collection of each sample, 200 μL of fresh medium was added to the well to maintain the culture media volume.

## ACKNOWLEDGEMENTS

We thank the Center for Instrumental Analysis and Metrology, Core Facility and Technical Support, and Center for Animal Experiment, Wuhan Institute of Virology, for providing technical assistance.

This work was supported by the National Key Research and Development Program of China (2018YFA0507204), the National Natural Sciences Foundation of China (31670165), Wuhan National Biosafety Laboratory, Chinese Academy of Sciences Advanced Customer Cultivation Project (2019ACCP-MS03), the Open Research Fund Program of the State Key Laboratory of Virology of China (2018IOV001).

